# Drug-Drug Interaction Extraction from Biomedical Text using Relation BioBERT with BLSTM

**DOI:** 10.1101/2022.08.31.506076

**Authors:** Maryam KafiKang, Abdeltawab Hendawi

## Abstract

Drug-drug interactions (DDIs) happen when two or more drugs interact. DDIs may change the effect of drugs in the body which can induce adverse effects and severe diseases for patients. As a result, detecting the interaction between drugs is essential. In the last years, many new DDIs have been found and added to the medical datasets with the rise in the number of discovered drugs. On the other hand, since a lot of this information is still in Biomedical articles and sources, there is a need for a method to extract DDIs information. However, despite the development of many techniques, attaining good prediction accuracy is the main issue. This paper proposes a deep learning approach that: 1) uses the power of Relation BioBERT (R-BioBERT) to detect and classify the DDIs and 2) employs the Bidirectional Long-Short Term Memory (BLSTM) to further increase the prediction quality. Not only does this paper study whether the two drugs have an interaction or not, but it also studies specific types of interactions between drugs. The paper also provides that using BLSTM can significantly increase the F-scores compared to the baseline model on the famous SemEval 2013, TAC 2018, and TAC 2019 DDI Extraction datasets.

## I Introduction

Drug-drug interactions (DDIs) occur when two or more drugs interact with each other, and their interactions impact the behavior of the drugs [1]. In some conditions, adverse drug reactions (ADRs) which are considered as health hazards and life-threatening issues may induce DDIs [2]. When a patient uses two or more drugs simultaneously, there is a risk of DDIs, which may endanger the patient’s health or even cause death. DDIs have become a bottleneck for drug administration, and patient safety [3]. As a result, DDIs are one of the most critical factors that affect drugs that cause side effects on patients’ health. Based on U.S centers’ reports, each year, ADRs kill 300 thousand people in the U.S. and Europe [4]. In addition, at most times, approximately 10% of people take five drugs or more simultaneously, and 20% of the elderly population takes at least ten drugs at the same time [5] which has dramatically increased the risk of ADRs. Since, DDIs provide vital information for patients, medical researchers, and doctors in pubHealth, doctors must have accurate knowledge about DDIs to prescribe drugs to patients.

With the rise in the usage of various drugs, it’s so important to have databases that store drug information. Keeping up-to-date all of these databases with the exponential growth of biomedical literature is a tough task [6], [7]. Some databases such as DrugBank [8], Therapeutic Target DB [9], and PharmGKB [10] are integrating to provide drug information including DDIs information for medical researchers and scientists. However, a considerable amount of DDIs information is still in biomedical articles instead of databases. Therefore, there is a need for automatic methods to extract DDIs from texts.

In recent years, the traditional methods for DDIs extraction like text mining [11] and statistical methods have been replaced by machine learning approaches [12], [13] which have shown better performances in DDIs extraction. However, the performance of the reported methods is still low, and there is still room for more improvements.

DDIs extraction tasks include discovering the drug name entities in the text and extracting interaction relations between drug entity pairs. In this work, we study three popular datasets that were introduced with SemEval 2013 [14], TAC 2018 [15], and TAC 2019 [16] DDIs Extraction challenge. We also incorporate the information from large-scale raw texts using a Bidirectional Encoder Representation From Transformers (BERT) [17] which is pre-trained on large-scale raw texts.

We develop a deep learning algorithm for DDIs extraction task based on Relation BioBERT (R-BioBERT) [18] and Bidirectional Long-Short Term Memory (BLSTM) [19]. We enrich the input sentence using the R-BioBERT model to extract the relation between drugs in a pair. Also, we employ the BLSTM model that takes the output of Relation-BioBERT as input and classifies the interaction in a pair into a specific DDI type. Using our deep learning model, we have achieved F1-macro 83.32% on SemEval 2013, 80.23% on TAC 2019, and 60.53% on TAC 2018 DDIs extraction showing that our model outperforms the baseline model (R-BioBERT).

The main contribution of this work can be summarized as follows:

- We develop a novel method that combines BLSTM with the Relation BioBERT to extract DDIs and classify their type of relations.
- The proposed model is evaluated on three datasets: SemEval 2013, TAC 2018, and TAC 2019 DDIs extraction.

Our results show that R-BioBERT with BLSTM (our method) outperforms the baseline model.

The rest of this paper is structured as follows: Section II describes the related works in the DDIs extraction task. Section III describes our proposed method, which is R-BioBERT with BLSTM. Section IV, the paper analyzes the experimental results and concludes the R-BioBERT with BLSTM with supported results.

The rest of this paper is structured as follows: Section II describes the related works in the DDIs extraction task. Section III describes our proposed method, which is R-BioBERT with BLSTM. Section IV, the paper analyzes the experimental results and concludes the R-BioBERT with BLSTM with supported results.

## II. Related works

DDIs extraction finds semantic relations between pairs of drugs. The main deep learning methods have been applied in the DDIs extraction task are supervised approaches [20]–[22]. Recently, recurrent neural networks (RNNs) [12], convolutional neural networks (CNNs) [13], and recursive neural networks (recursive NNs) [23] in DDIs extraction have shown that they can capture meaningful information and outperform traditional machine learning methods.

CNNs is a robust deep learning method that draws great attention in many real-world applications, including image classification [24], object detection [25], and many engineering applications [26]. CNNs can also be applied in natural language processing tasks like sentiment analysis [27], search query [28], and semantic parsing [29]. CNNs have been applied in DDIs extraction tasks. The first application of CNNs in the DDIs extraction task has developed by Liu, Shengyu, et al.[30]. Asada et al.[31] designed a method that is a combination of attention mechanisms with CNN that outperformed the CNN-based model [30] in DDIs extraction task. Some works like [32], [33] applied deep CNN architecture by increasing the depth of networks. In addition, Asada, Masaki, et al.[9] proposed a model that applied CNNs with a graph that encoded the sentences and molecular drug pair in the DDIs extraction task. The MCCNN [34] proposed a method based on distributed word embedding and a multichannel convolutional neural network for biomedical relation extraction. Sun et al.[35] developed a method based on BLSTM to learn the semantic knowledge from texts and CNN to achieve sentence features.

RNNs are another popular deep learning methods that are great at capturing sentence features. In comparison to CNNs, RNNs are more suitable for NLP tasks. Kavuluru et al.[36] developed a method by combining an original word-based RNN with a character-based RNN. D.Huang et al.[37] proposed a two-stage LSTM model which is composed of a SVM model to recognize the negative and positive DDIs and a LSTM model to classify DDIs into a specific category. Z.Yi, S.Li et al.[38] proposed 2ATT-RNN, which has two attention layers, the word level attention layer and the sentence layer attention layer. Another RNNs-base method is the joint AB-LSTM model, which is proposed by [22] in the DDIs extraction task. The paper [39] presented a position-aware attention mechanism in their model (PM-BLSTM) that combines the relative position information of the target entities with the hidden states of the BiLSTM layer. Although these deep learning approaches have achieved great results, there are possibilities for more improvement.

Recently, the application of contextualized embedding-based methods has increased, and these methods have drawn great attention [40]. Deep transformers-based methods have been trained by contextualized embedding-based methods [40] on a large text data. For instance, in many NLP tasks, BERT [17], a pre-trained language model, has been applied and attracting great attention. BERT takes advantage of a bidirectional encoder transformer to capture richer context than other word embedding methods like Glove [41], and Word2vec [42]. Consequently, BERT boosts the model’s performance by integrating contextual information in sentences. For instance, Datta et al.[43] developed a BERT based model to extract DDIs in the sentences. In addition, Zaikis et al.[44] proposed a deep learning model based on Transformer [45] architecture and the BERT language model for DDIs extraction task. Based on this, some bio-specific BERT models which are trained with large-scale biomedical corpora, like BioBERT [46], SciBERT [10], and Med-BERT [47] have been applied in several DDIs extraction tasks. For example, [48], [49] combine BioBERT and SciBERT to obtain richer sequence semantic information.

## III. Method

We propose a novel method based on R-BioBERT and BLSTM for the DDIs extraction task. We illustrate the overview of the proposed DDIs extraction model in Figure 1. The model extracts interactions from a sentence and classifies them into a specific DDIs type. In this section, we first present BERT, BioBERT, and R-BioBERT algorithms and then introduce our proposed model (R-BioBERT with BLSTM).

**Fig. 1:**
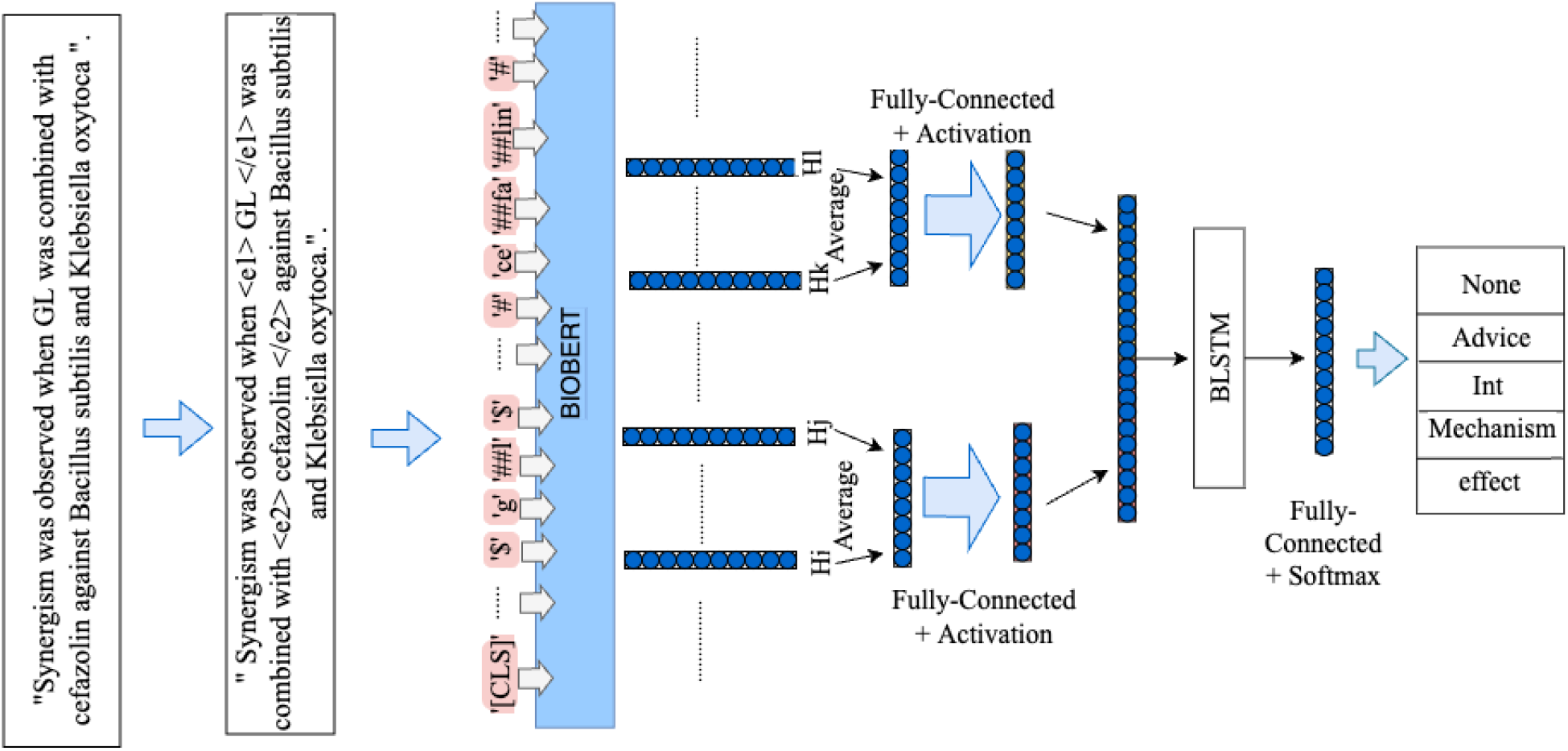
The model Architecture.

### A. BERT language model

We need language representation models in NLP tasks to learn word representations from unlabeled texts. Some previous language models like Glove and Word2Vec are context-free, which means they focus on learning word representation without considering the context of words in sentences. Recent language representation models like ELMO [50] and Cove [51] are context-sensitive.

BERT is a pre-trained language representation model that stands for bidirectional encoder representation from transformers. BERT utilizes a context-sensitive word representation model [17]. BERT, a general-purpose language model, has been trained on large-scale corpora, including English Wikipedia and Books Corpus, to obtain contextualized representations of each word in sentences. BERT uses the encoder part of Transformers to encode a text’s semantic and syntactic information in the embedding form, which is a language model. The procedure of pre-training of BERT has two objectives: masked language model (MLM) and next sentence prediction (NSP).

The MLM randomly masks some of the tokens from input and sets the optimization objective to predict the original vocabulary id of the masked word according to its context. In the NSP, The BERT model has been trained to predict the text-pair representation. In BERT, a special classification token ‘[CLS]’ is added to every input sequence, including a single or two sentences *< Question, Answer >*. For classification tasks, the final hidden state corresponds to the ‘[CLS]’ tokens used as an aggregate sequence representation [17].

### B. BioBERT

BioBERT [17] stands for Bidirectional Encoder Representations from Transformers for Biomedical Text Mining. BioBERT is based on BERT, a contextualized language representation model trained on various general and biomedical datasets, including PubMed Abstracts and PMC Fulltext articles. BioBERT has outperformed BERT and stateof-the-art models in many NLP tasks like Named Entity Recognition (NER) from Bio-medical data, relation extraction, and question & answer in the biomedical field.

Biomedical domain-specific literature includes specific nouns and terms that are not in general corpora. Consequently, using general-purpose language representation models like BERT for NLP tasks in the Biomedical domain can not achieve a good performance. To overcome this problem, this work uses BioBERT, which is a biomedical domain-specific Language Representation Model based on the BERT.

### C. Relation BERT

The goal of relation classification tasks is to predict semantic relations between pairs of nominals. For instance, if we have a sentence s and a pair of two nominals which are *e*_1_ and *e*_2_, the object is to identify the relation between *e*_1_ and *e*_2_ [52]. Many deep learning approaches have been applied in relation classification tasks [53], [54] and most of them are based on convolutional or recurrent neural networks. Recently, the pre-trained BERT model has been applied to multiple NLP tasks and obtained state-of-the-art results on classification and SQuAD question answering problems [55]. In classification problems, the information of the sentence is important, although in Relation classification tasks besides the information of the sentence, the information of the target entities is important as well.

For instance, Wu S, et al.[18] applied the pre-trained BERT model to relation classification. The paper leverages the pretrain BERT language model and incorporates information from the target entities in the sentence to tackle the relation classification task. In general, R-BERT consists of two components: the pre-trained BERT as the feature representation and additional layers as the relation classifier.

The major distinction between the BERT and the Relation BERT is that BERT uses the final hidden state vectors of ‘[CLS]’ as the input for the classification layer although RBERT uses the final hidden state vectors of ‘[CLS]’ and two entities of interest as the inputs for their classification layers. ‘[CLS]’ as the first token of each sentence is appended to the beginning of each sentence. The final hidden state from the transformer output corresponding to the first token which is ‘[CLS]’ is used as the sentence representation for classification tasks.

### D. Model Architecture

We propose a method based on Relation BioBERT and BLSTM. According to the architecture of the R-BERT model, the location of drugs that are in interaction should be detected and a masked symbol should be added before and after each target drug. In our model, since drugs do not have a fixed length, we add *< e*1 *>* before the name of the first drug and add *< /e*1 *>* at the end of the first drug. We do the same for the second drug but we replace *< e*1 *>* with *< e*2 *>*. After detecting the location of the first and the second drugs that are in an interaction, we feed input to BioBERT for generating the feature representation.

For example, after insertion of the special separate tokens, for a sentence with target entities ***”Ganoderma lucidum extract”*** and ***”antibiotics”*** will become to: *”Antimicrobial activity of < e*1 *>* ***Ganoderma lucidum extract*** *< /e*1 *> alone and in combination with some < e*2 *>* ***antibiotics*** *< /e*2 *>*.*”*

In our model, we use BioBERT instead of BERT, because the DDIs extraction task is a biomedical relation extraction task, and our model needs a language model that has been trained on Biomedical corpora instead of only general corpora. Given a sentence *S* with entity *e*_1_ and *e*_2_, suppose its final hidden state output from BioBERT module is *H*. Suppose vectors *H*_*i*_ to *H*_*j*_ are the final hidden state vectors from BioBERT for entity *e*_1_, and *H*_*k*_ to *H*_*m*_ are the final hidden state vectors from BioBERT for entity *e*_2_. We apply the average operation to get a vector representation for the two target entities. Then after an activation operation (i.e. tanh), we add a fully connected layer to each average vectors, and the output for *e*_1_ and *e*_2_ are 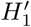 and 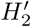 respectively.

We concatenate 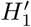 and 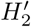 and apply the BLSTM layer to each separately. Then, we pass the output of BLSTM through Fully-connected layer and a softmax layer. The architecture of the proposed model is illustrated in Figure 1. The reader can find more information about Relation BERT in [18].

## IV. Experimental evaluation

This section explains the experimental settings, drug databases, data preprocessing steps, and final results. Then we compare our results with the state-of-the-art models.

### A. Experimental setup

Table I shows the major parameters used in our experiments. The experiments are conducted on a computer with a Windows operating system with a single Nvidia GeForce RTX 2070 with 8GB memory. We implemented our model with the Pytorch library and the Python programming language.

**TABLE I:** Parameter settings

|  |  |
| --- | --- |
| Batch size | 8 |
| Max sentence length | 400 |
| Adam learning rate | 2e-5 |
| Number of epochs | 5 |
| Dropout rate | 0.1 |

### B. Datasets

We utilize three datasets which are the DDIs corpus from SemEval 2013 [14], TAC 2018 [15], and TAC 2019 DDIs extraction [16] in our work.

#### 1. SemEval 2013 DDIs extraction

The DDIs Extraction 2011 [56] was designed to be used in detection tasks of DDIs biomedical texts. The DDIs Extraction 2013 followed up on the DDIs Extraction 2011 as a new edition to support more tasks instead of DDIs like recognition and classification of pharmacological substances [14]. For extraction of DDIs task, this dataset consists of DrugBank with 730 documents and MEDLINE with 175 abstracts. For training and evaluation of different systems, the dataset is divided into training with 714 documents and test with 191 documents [14]. This dataset contains four effective DDI types: Advice, Effect, Int, and Mechanism. The relation between drug entity pairs that do not belong to any four DDIs mentioned before is determined as negative and significantly more than positive DDI instances.

- Advice: This is referred to as advice or recommendation about the simultaneous use of two drugs in the document; for example, “Caution should be used when alosetron and ketoconazole are administered concomitantly.”
- Effect: This is referred to as an effect of the DDI or pharmacodynamic mechanism of interaction. For example, “PGF2alpha produced significantly increased vasoconstriction after a single administration of oxytocin.”
- Int: This is referred to as an interaction between drugs without providing more information. For instance, “Clinical implications of warfarin interactions with five sedatives.”
- Mechanism: This is referred to as a pharmacokinetic mechanism description, as in “Withdrawal of rifampin decreased the warfarin requirement by 50%.”.
- Negative: There is no interaction between two drug entity pairs. For example, “Ibogaine, but not 18-MC, decreases heart rate at high doses.”

#### 2) TAC 2018 and TAC 2019 DDIs extraction

The U.S. Food and Drug Administration (FDA) and the National Library of Medicine (NLM) worked together to prepare a dataset to be effective for the deployment of drug safety information [15]. The main goal of the TAC track is to evaluate NLP approaches for information extraction performance in DDIs. In addition, TAC provides data for other tasks such as Extraction entities and relations, and Normalization [15]. The TAC 2018 DDIs track dataset includes 325 structured product labels (SPLs), are divided into a training set with 22 drug labels and a test set with 57 drug labels. The type of DDIs is divided into three groups:

- Pharmacokinetic (PK)
- Pharmacodynamic (PD)
- Unspecified (U)

The difference between TAC 2019 and TAC 2018 DDI extraction is that TAC 2018 includes the information from SPLs and other text types like literature and social media. TAC 2019 DDIs Extraction dataset includes 406 structured product labels (SPLs), which consist of the training set with 211 drug labels, and the test set with 81 drug labels. However, the type of DDIs is the same as DDIs in TAC 2018 DDI extraction. The statistics of the dataset with the official data split are shown in Table II.

**TABLE II:** Statistics of TAC 2018 and TAC 2019 dataset

| DDIs | Train |  | Test |  |
| --- | --- | --- | --- | --- |
|  | TAC 2018 | TAC 2019 | TAC 2018 | TAC 2019 |
| Pharmacodynamic | 47 | 553 | 335 | 292 |
| Pharmacokinetic | 60 | 494 | 296 | 118 |
| Unspecified | 62 | 665 | 440 | 202 |

### C. Data preprocessing

For TAC 2018, and TAC 2019 DDIs extraction datasets, the following steps are applied:

- Since a drug does not interact with itself [57], we remove instances with the the same drug names in a pair. We also remove instances that have only one drug in a sentence.
- We add special token *< e*1 *>* before and *< /e*1 *>* after the first drug and add *< e*2 *>* before and *< /e*2 *>* after the second drug to detect the location of two drugs in a pair. Unlike most related studies, we keep the original name of drugs.

In semEval 2013 DDIs extraction, the number of negative types of interactions is significantly more than that of positive interactions. This means that the dataset suffers from an imbalance class distribution problem which decreases the performance accuracy of the deep learning model [58]. To alleviate the imbalance problem, Zhao, Zhehuan, et al.[21] constructed a less imbalanced corpus by applying some rules to remove extra negative instances from SemEval 2013 DDIs Extraction dataset. We use almost the same number of data pairs from this study’s released code and data. The only modification we add to this dataset is applying the second step of the TAC preprocessing method to that.

### D. Evaluation Metrics

F1-Macro and F1-weighted are the primary evaluation metrics that are widely used in the DDIs extraction task. In this study, we also evaluate our model (R-BioBERT with BLSTM) performance with weighted-average and macro-average F1score on all types for SemEval 2013 and macro-average on all types for TAC 2018 and TAC 2019 DDIs extractions datasets:

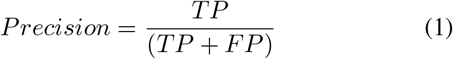

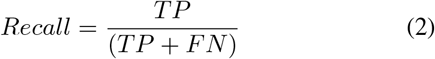

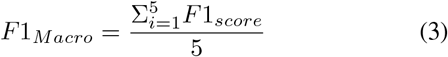

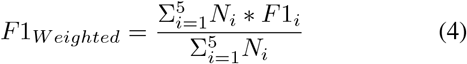

Where *N*_*i*_ is the number of instances in class i, TP (true positive) represents the number of correctly classified positive instances, FP (false positive) is the number of negative instances which are misclassified as positive instances, and FN (false negative) is the number of positive instances that are misclassified as negative. Precision is the ratio of correctly predicted positive observations to the total predicted observations. Recall is the ratio of correctly predicted positive observations to all observations in the actual class. The F1-score metric is the harmonic mean of Precision and Recall metrics. The F1 macro-averaged score or F1-macro is an unweighted mean of all the per-class F1 scores, which treats all classes equally. The weighted-average F1 score or F1-weighted is calculated by taking the mean of all per-class F1 scores while considering the number of actual samples of each class.

### E. Results

#### 1) Results on SemEval 2013 DDIs extraction

Table III shows the performance of our proposed model and the state-ofthe-art models on the SemEval 2013 DDIs extraction task to determine our work’s position. The macro-average F1-score of the proposed model is calculated based on **five classes** including **Negative DDI** class. To compare the performance of our model with related studies, we mentions results of some DDIs extraction tasks that applied RNNs or CNNs or BERT including MCCNN [34], RHCNN [35], join AB-LSTM [22], TP-DDI [44], BERT-D2, BERTChem [48], [43] and RBERT(the baseline method) [59]. In the table III we can see that the BERT-based model achieved a higher overall f1-score compared to the RNNs and CNNs-based models.

**TABLE III:**
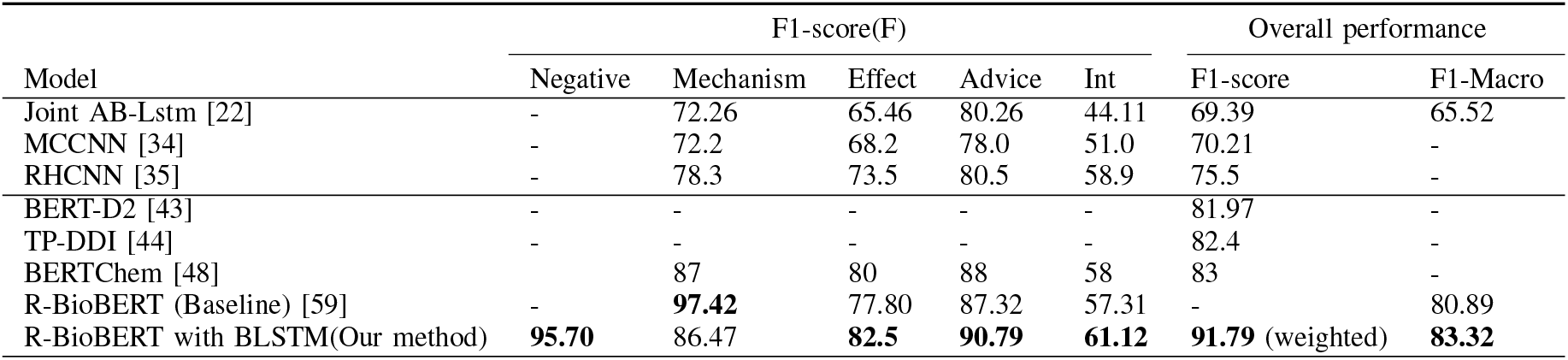
Model performance on SemEval 2013 DDI extraction

| Model | F1-score(F) |  |  |  |  | Overall performance |  |
| --- | --- | --- | --- | --- | --- | --- | --- |
|  | Negative | Mechanism | Effect | Advice | Int | F1-score | F1-Macro |
| Joint AB-Lstm [22] | - | 72.26 | 65.46 | 80.26 | 44.11 | 69.39 | 65.52 |
| MCCNN [34] | - | 72.2 | 68.2 | 78.0 | 51.0 | 70.21 | - |
| RHCNN [35] | - | 78.3 | 73.5 | 80.5 | 58.9 | 75.5 | - |
| BERT-D2 [43] | - | - | - | - | - | 81.97 | - |
| TP-DDI [44] | - | - | - | - | - | 82.4 | - |
| BERTChem [48] | - | 87 | 80 | 88 | 58 | 83 | - |
| R-BioBERT (Baseline) [59] | - | **97.42** | 77.80 | 87.32 | 57.31 | - | 80.89 |
| R-BioBERT with BLSTM(Our method) | **95.70** | 86.47 | **82.5** | **90.79** | **61.12** | **91.79** (weighted) | **83.32** |

Table III also shows that in CNNs and LSTM-based models, the Joint AB-Lstm model has the lowest overall F1-score (69.39%) and the best F1-score for this model belongs to Mechanism (72.26%), and its worst performance belong to Int (44.11%). MCCNN has a better overall F1-score of 70.21% in comparison to the Joint AB-Lstm, although it is less than RHCNN (75.5%).

In BERT-based models, BERT-D2 has the lowest overall F1 score (81.97%), and it didn’t report the model’s performance on each DDIs. TP-DDI model has a better overall f1-score (82.4%) in comparison to BERT-D2. However, its overall F1score is less than BERTChem (83%). The R-BioBERT, which is our baseline model, achieved overall F1-Macro (80.89%). R-BioBERT with BLSTM (our model) achieves the best overall weighted-average F1 (91.79) and F1-Macro (83.32) in comparison to the state-of-the-art models.

R-BioBERT has the highest F1-score (97.42%), and MCCNN has the lowest F1-score (72.2 %) in Mechanism. The highest F1-score in Negative (95.70%), Effect (82.5%), Advice (90.79%), and Int (61.12%) belongs to the R-BioBERT with BLSTM (our model). In addition, the worst F1-score for Effect and Int belongs to the Joint AB-Lstm. In Advice, The highest F1 score belongs to the R-BioBERT with BLSTM (our model), and the worst F1 score belongs MCCNN. The highest scores are bolded in table III.

The highest F1-score (91.79%) and F1-macro (83.32%) in overall performance belong to the R-BioBERT with the BLSTM model.

Our proposed method, R-BioBERT with BLSTM, significantly beats previous CNN and LSTM-based models, especially the baseline method [59]. The F1 macro of R-BioBERT with BLSTM is 83.32, which is much better than the previous best solution on SemEval 2013 dataset. Our model also achieve better performance with distinction, in case of the **Negative, Effect, Int** classes, as compared to previous researches. In all research, the Int class has limited performance regardless to model architecture. The low performance in the Int class is likely due to insufficient training data.

#### 2) Results on TAC 2018

Table IV shows the performance of our proposed model and the state-of-the-art model on the TAC 2018 DDIs extraction task. For the TAC 2018 DDIs extraction, F1-macro of proposed model is calculated based on three classes. Table IV shows that Tang et al.[60] has the lowest F1 score (40.90%), although it used NLM-180 and HS datasets besides TAC 2018. The model proposed in [61] has a better F1-score (56.98%) compare to [60]. R-BioBERT with BLSTM achieves the best F1-macro score in comparison to the state-of-the-art models. One reason for the low F1-score can be the lack of data in TAC 2018. This dataset is much smaller than SemEval 2013 and TAC 2019. So it is not surprising to see the relatively small F1-score for TAC 2018 model.

**TABLE IV:** Model performance on TAC 2018 DDI extraction

| Model | F1-score(F) |  |  | Overall performance |
| --- | --- | --- | --- | --- |
|  | Unspecified | Pharmacokinetic (PK) | Pharmacodynamic (Specific) | F1-score |
| [60] | - | - | - | 40.90 |
| [61] | - | - | - | 56.98 |
| R-BioBERT with BLSTM(Our method) | 64 | 58.8 | 58.8 | **60.53** (Macro) |

##### 3) Results on TAC 2019

Table V shows the performance of our proposed model and the state-of-the-art model on the TAC 2019 DDIs extraction task. For the TAC 2019 DDIs extraction, The macro-average F1-score of the proposed model is calculated based on three classes. In the table V we can see that Mahajan et al.[62] has the worst f1-score (40.39%). Although UTDHLTRI 2 [16] has better f1-score (49.2%), it is less than the F1-score of IBMResearch 1 [16] (50.1%). R-BioBERT with BLSTM achieves the highest F1-score, 80.26%. Our proposed model (R-BioBERT with BLSTM) outperforms the state-of-the-art models by a large margin.

**TABLE 5:** Model performance on TAC 2019 DDI extraction

| Overall performance | F1-score(F) |  |  |  |
| --- | --- | --- | --- | --- |
|  | Unspecified | Pharmacokinetic (PK) | Pharmacodynamic (Specific) | F1-score |
| [62] | 48.3 | 63.2 | 43.4 | 40.39 |
| UTDHLTRI 2 [16] | - | - | - | 49.2 |
| IBMResearch 1 [16] | - | - | - | 50.1 |
| R-BioBERT with BLSTM(Our method) | 76.2 | 81.2 | 83.3 | **80.23** (Macro) |

## V Conclusion

In this work, we proposed a method based on Relation-BioBERT and BLSTM for the DDI extraction. Our results show that adding BLSTM to R-BioBERT can increase the F1score by **3**% compared to the baseline model, which doesn’t contain BLSTM. We also show that our model outperforms the state-of-the-art models in DDI extraction on Semeval 2013, TAC 2018, and TAC 2019 DDI extraction. Our model achieves F1-macro 83.32% on SemEval, 80.23% on TAC 2018, and 60.53% on TAC 2018 DDIs extraction.

## References

[1] V. Miranda, A. Fede, M. Nobuo, et al., “Adverse drug reactions and drug interactions as causes of hospital admission in oncology,” Journal of pain and symptom management, vol. 42, no. 3, pp. 342–353, 2011.

[2] J. Lazarou, B. H. Pomeranz, and P. N. Corey, “Incidence of adverse drug reactions in hospitalized patients: A meta-analysis of prospective studies,” Jama, vol. 279, no. 15, pp. 1200–1205, 1998.

[3] M. L. Becker, M. Kallewaard, P. W. Caspers, L. E. Visser, H. G. Leufkens, and B. H. Stricker, “Hospitalisations and emergency department visits due to drug–drug interactions: A literature review,” Pharmacoepidemiology and drug safety, vol. 16, no. 6, pp. 641–651, 2007.

[4] R. Businaro, “Why we need an efficient and careful pharmacovigilance?” Journal of pharmacovigilance, 2013.

[5] C. M. Hohl, J. Dankoff, A. Colacone, and M. Afilalo, “Polypharmacy, adverse drug-related events, and potential adverse drug interactions in elderly patients presenting to an emergency department,” Annals of emergency medicine, vol. 38, no. 6, pp. 666–671, 2001.

[6] R. P. Paczynski, G. C. Alexander, V. M. Chinchilli, and S. P. Kruszewski, “Quality of evidence in drug compendia supporting off-label use of typical and atypical antipsychotic medications,” International Journal of Risk & Safety in Medicine, vol. 24, no. 3, pp. 137–146, 2012.

[7] A. Rodriguez-Terol, M. Caraballo, D. Palma, et al., “Quality of interaction database management systems,” Farmacia Hospitalaria (English Edition), vol. 33, no. 3, pp. 134–146, 2009.

[8] W. Ammar, D. Groeneveld, C. Bhagavatula, et al., “Construction of the literature graph in semantic scholar,” arXiv preprint 1805.02262, 2018.

[9] M. Asada, M. Miwa, and Y. Sasaki, “Enhancing drug-drug interaction extraction from texts by molecular structure information,” arXiv preprint 1805.05593, 2018.

[10] I. Beltagy, K. Lo, and A. Cohan, “Scibert: A pretrained language model for scientific text,” arXiv preprint 1903.10676, 2019.

[11] S. Yan, X. Jiang, and Y. Chen, “Text mining driven drug-drug interaction detection,” in 2013 IEEE international conference on bioinformatics and biomedicine, IEEE, 2013, pp. 349–354.

[12] R. J. Williams and D. Zipser, “A learning algorithm for continually running fully recurrent neural networks,” Neural computation, vol. 1, no. 2, pp. 270–280, 1989.

[13] Y. LeCun, L. Bottou, Y. Bengio, and P. Haffner, “Gradient-based learning applied to document recognition,” Proceedings of the IEEE, vol. 86, no. 11, pp. 2278–2324, 1998.

[14] M. Herrero-Zazo, I. Segura-Bedmar, P. Martinez, and T. Declerck, “The ddi corpus: An annotated corpus with pharmacological substances and drug–drug interactions,” Journal of biomedical informatics, vol. 46, no. 5, pp. 914–920, 2013.

[15] D. Demner-Fushman, K. W. Fung, P. Do, R. D. Boyce, and T. R. Goodwin, “Overview of the tac 2018 drugdrug interaction extraction from drug labels track.,” in TAC, 2018.

[16] T. R. Goodwin, D. Demner-Fushman, K. W. Fung, and P. Do, “Overview of the tac 2019 track on drug-drug interaction extraction from drug labels.,” in TAC, 2019.

[17] J. Devlin, M.-W. Chang, K. Lee, and K. Toutanova, “Bert: Pre-training of deep bidirectional transformers for language understanding,” arXiv preprint 1810.04805, 2018.

[18] S. Wu and Y. He, “Enriching pre-trained language model with entity information for relation classification,” in Proceedings of the 28th ACM international conference on information and knowledge management, 2019, pp. 2361–2364.

[19] M. Schuster and K. K. Paliwal, “Bidirectional recurrent neural networks,” IEEE transactions on Signal Processing, vol. 45, no. 11, pp. 2673–2681, 1997.

[20] L. Hong, J. Lin, S. Li, et al., “A novel machine learning framework for automated biomedical relation extraction from large-scale literature repositories,” Nature Machine Intelligence, vol. 2, no. 6, pp. 347–355, 2020.

[21] Z. Zhao, Z. Yang, L. Luo, H. Lin, and J. Wang, “Drug drug interaction extraction from biomedical literature using syntax convolutional neural network,” Bioinformatics, vol. 32, no. 22, pp. 3444–3453, 2016.

[22] S. K. Sahu and A. Anand, “Drug-drug interaction extraction from biomedical texts using long short-term memory network,” Journal of biomedical informatics, vol. 86, pp. 15–24, 2018.

[23] R. Socher, C. C.-Y. Lin, A. Y. Ng, and C. D. Manning, “Parsing natural scenes and natural language with recursive neural networks,” in ICML, 2011.

[24] Y. Jiang, L. Chen, H. Zhang, and X. Xiao, “Breast cancer histopathological image classification using convolutional neural networks with small se-resnet module,” PloS one, vol. 14, no. 3, e0214587, 2019.

[25] X. Dai, Y. Chen, B. Xiao, et al., “Dynamic head: Unifying object detection heads with attentions,” in Proceedings of the IEEE/CVF Conference on Computer Vision and Pattern Recognition, 2021, pp. 7373–7382.

[26] M. M. Behzadi and H. T. Ilieş, “Real-time topology optimization in 3d via deep transfer learning,” Computer-Aided Design, vol. 135, p. 103 014, 2021.

[27] M. E. Basiri, S. Nemati, M. Abdar, E. Cambria, and U. R. Acharya, “Abcdm: An attention-based bidirectional cnn-rnn deep model for sentiment analysis,” Future Generation Computer Systems, vol. 115, pp. 279– 294, 2021.

[28] Y. Shen, X. He, J. Gao, L. Deng, and G. Mesnil, “Learning semantic representations using convolutional neural networks for web search,” in Proceedings of the 23rd international conference on world wide web, 2014, pp. 373–374.

[29] W.-t. Yih, X. He, and C. Meek, “Semantic parsing for single-relation question answering,” in Proceedings of the 52nd Annual Meeting of the Association for Computational Linguistics (Volume 2: Short Papers), 2014, pp. 643–648.

[30] S. Liu, B. Tang, Q. Chen, and X. Wang, “Drug-drug interaction extraction via convolutional neural networks,” Computational and mathematical methods in medicine, vol. 2016, 2016.

[31] M. Asada, M. Miwa, and Y. Sasaki, “Extracting drugdrug interactions with attention CNNs,” in BioNLP 2017, Vancouver, Canada, Association for Computational Linguistics, Aug. 2017, pp. 9–18. DOI: 10.18653/v1/W17-2302. [Online]. Available: https://aclanthology.org/W17-2302.

[32] I. N. Dewi, S. Dong, and J. Hu, “Drug-drug interaction relation extraction with deep convolutional neural networks,” in 2017 IEEE International Conference on Bioinformatics and Biomedicine (BIBM), IEEE Computer Society, 2017, pp. 1795–1802.

[33] X. Sun, L. Ma, X. Du, J. Feng, and K. Dong, “Deep convolution neural networks for drug-drug interaction extraction,” in 2018 ieee international conference on bioinformatics and biomedicine (bibm), IEEE, 2018, pp. 1662–1668.

[34] C. Quan, L. Hua, X. Sun, and W. Bai, “Multichannel convolutional neural network for biological relation extraction,” BioMed research international, vol. 2016, 2016.

[35] X. Sun, K. Dong, L. Ma, et al., “Drug-drug interaction extraction via recurrent hybrid convolutional neural networks with an improved focal loss,” Entropy, vol. 21, no. 1, p. 37, 2019.

[36] R. Kavuluru, A. Rios, and T. Tran, “Extracting drugdrug interactions with word and character-level recurrent neural networks,” in 2017 IEEE International Conference on Healthcare Informatics (ICHI), IEEE, 2017, pp. 5–12.

[37] D. Huang, Z. Jiang, L. Zou, and L. Li, “Drug–drug interaction extraction from biomedical literature using support vector machine and long short term memory networks,” Information sciences, vol. 415, pp. 100–109, 2017.

[38] Z. Yi, S. Li, J. Yu, et al., “Drug-drug interaction extraction via recurrent neural network with multiple attention layers,” in International Conference on Advanced Data Mining and Applications, Springer, 2017, pp. 554–566.

[39] D. Zhou, L. Miao, and Y. He, “Position-aware deep multi-task learning for drug–drug interaction extraction,” Artificial intelligence in medicine, vol. 87, pp. 1–8, 2018.

[40] Y. Peng, S. Yan, and Z. Lu, “Transfer learning in biomedical natural language processing: An evaluation of bert and elmo on ten benchmarking datasets,” arXiv preprint 1906.05474, 2019.

[41] J. Pennington, R. Socher, and C. D. Manning, “Glove: Global vectors for word representation,” in Proceedings of the 2014 conference on empirical methods in natural language processing (EMNLP), 2014, pp. 1532–1543.

[42] T. Mikolov, K. Chen, G. Corrado, and J. Dean, “Efficient estimation of word representations in vector space,” arXiv preprint 1301.3781, 2013.

[43] T. T. Datta, P. C. Shill, and Z. Al Nazi, “Bert-d2: Drugdrug interaction extraction using bert,” in 2022 International Conference for Advancement in Technology (ICONAT), IEEE, 2022, pp. 1–6.

[44] D. Zaikis and I. Vlahavas, “Tp-ddi: Transformer-based pipeline for the extraction of drug-drug interactions,” Artificial Intelligence in Medicine, vol. 119, p. 102 153, 2021.

[45] T. Wolf, L. Debut, V. Sanh, et al., “Transformers: State-of-the-art natural language processing,” in Proceedings of the 2020 Conference on Empirical Methods in Natural Language Processing: System Demonstrations, Online: Association for Computational Linguistics, Oct. 2020, pp. 38–45. DOI: 10.18653/v1/2020.emnlp-demos.6. [Online]. Available: https://aclanthology.org/2020.emnlp-demos.6.

[46] J. Lee, W. Yoon, S. Kim, et al., “Biobert: A pretrained biomedical language representation model for biomedical text mining,” Bioinformatics, vol. 36, no. 4, pp. 1234–1240, 2020.

[47] L. Rasmy, Y. Xiang, Z. Xie, C. Tao, and D. Zhi, “Medbert: Pretrained contextualized embeddings on largescale structured electronic health records for disease prediction,” NPJ digital medicine, vol. 4, no. 1, pp. 1–13, 2021.

[48] I. Mondal, “Bertchem-ddi: Improved drug-drug interaction prediction from text using chemical structure information,” arXiv preprint 2012.11599, 2020.

[49] M. Asada, M. Miwa, and Y. Sasaki, “Using drug descriptions and molecular structures for drug–drug interaction extraction from literature,” Bioinformatics, vol. 37, no. 12, pp. 1739–1746, 2021.

[50] M. E. Peters, M. Neumann, M. Iyyer, et al., “Deep contextualized word representations,” in Proceedings of the 2018 Conference of the North American Chapter of the Association for Computational Linguistics: Human Language Technologies, Volume 1 (Long Papers), New Orleans, Louisiana: Association for Computational Linguistics, Jun. 2018, pp. 2227–2237. DOI: 10.18653/v1/N18-1202. [Online]. Available: https://aclanthology.org/N18-1202.

[51] B. McCann, J. Bradbury, C. Xiong, and R. Socher, “Learned in translation: Contextualized word vectors,” Advances in neural information processing systems, vol. 30, 2017.

[52] I. Hendrickx, S. N. Kim, Z. Kozareva, et al., “Semeval-2010 task 8: Multi-way classification of semantic relations between pairs of nominals,” arXiv preprint 1911.10422, 2019.

[53] C. Nogueira dos Santos, B. Xiang, and B. Zhou, “Classifying relations by ranking with convolutional neural networks,” arXiv e-prints, arXiv–1504, 2015.

[54] J. Lee, S. Seo, and Y. S. Choi, “Semantic relation classification via bidirectional lstm networks with entity-aware attention using latent entity typing,” Symmetry, vol. 11, no. 6, p. 785, 2019.

[55] P. Rajpurkar, J. Zhang, K. Lopyrev, and P. Liang, “Squad: 100,000+ questions for machine comprehension of text,” arXiv preprint 1606.05250, 2016.

[56] I. Segura-Bedmar, P. Martinez Fernández, and D. Sánchez Cisneros, “The 1st ddiextraction-2011 challenge task: Extraction of drug-drug interactions from biomedical texts,” 2011.

[57] M. F. M. Chowdhury and A. Lavelli, “Fbk-irst: A multiphase kernel based approach for drug-drug interaction detection and classification that exploits linguistic information,” in Second Joint Conference on Lexical and Computational Semantics (* SEM), Volume 2: Proceedings of the Seventh International Workshop on Semantic Evaluation (SemEval 2013), 2013, pp. 351–355.

[58] Y. Sun, A. K. Wong, and M. S. Kamel, “Classification of imbalanced data: A review,” International journal of pattern recognition and artificial intelligence, vol. 23, no. 04, pp. 687–719, 2009.

[59] D. P. Nguyen and T. B. Ho, “Drug-drug interaction extraction from biomedical texts via relation bert,” in 2020 RIVF International Conference on Computing and Communication Technologies (RIVF), IEEE, 2020, pp. 1–7.

[60] S. Tang, Q. Zhang, T. Zheng, et al., “Two step joint model for drug drug interaction extraction,” arXiv preprint 2008.12704, 2020.

[61] T. Tran, R. Kavuluru, and H. Kilicoglu, “A multitask learning framework for extracting drugs and their interactions from drug labels,” arXiv preprint 1905.07464, 2019.

[62] D. Mahajan, A. Poddar, and Y.-T. Lin, “A hybrid model for drug-drug interaction extraction from structured product labeling documents,” in TAC, 2019.

